# Non-invasive measurement of hepatic venous oxygen saturation (ShvO_2_) with quantitative susceptibility mapping in normal mouse liver and livers bearing colorectal metastases

**DOI:** 10.1101/193656

**Authors:** E. Finnerty, R. Ramasawmy, J. O’Callaghan, J. Connell, M. F. Lythgoe, K. Shmueli, D. Thomas, S. Walker-Samuel

## Abstract

**Purpose:** The purpose of this prospective study was to investigate the potential of QSM to non-invasively measure hepatic venous oxygen saturation (ShvO_2_).

**Materials & Methods:** All animal studies were performed in accordance with the UK Home Office Animals Science Procedures Act (1986) and UK National Cancer Research Institute (NCRI) guidelines. QSM data was acquired from a cohort of mice (n=10) under both normoxic (medical air, 21% O2/ balance N), and hyperoxic conditions (100% O_2_). Susceptibility measurements were taken from large branches of the portal and hepatic vein under each condition and were used to calculate venous oxygen saturation in each vessel. Blood was extracted from the IVC of three mice under norm- and hyperoxic conditions, and oxygen saturation was measured using a blood gas analyser to act as a gold standard. QSM data was also acquired from a cohort of mice bearing colorectal liver metastases (CRLM). SvO_2_ was calculated from susceptibility measurements made in the portal and hepatic veins, and compared to the healthy animals.

**Results:** SvO_2_ calculated from QSM measurements showed a significant increase of 14.93% in the portal vein (p < 0.05), and an increase of 21.39% in the hepatic vein (p < 0.01). Calculated results showed excellent agreement with those from the blood gas analyser (26.14% increase). ShvO_2_ was significantly lower in the disease cohort (30.18 ± 11.6%), than the healthy animals (52.67 ± 17.8%) (p < 0.05), but differences in the portal vein were not significant.

**Conclusion:** QSM is a feasible tool for non-invasively measuring hepatic venous oxygen saturation and can detect differences in oxygen consumption in livers bearing colorectal metastases.

## Introduction

The measurement of venous blood susceptibility and haemoglobin oxygen saturation (SvO_2_) using Quantitative Susceptibility Mapping (QSM) has been the focus of several studies in recent years^[1-4]^. It has been shown in both animal models^[3]^, and humans^[2, 5, 6]^ that QSM can quantify changes in deoxyhaemoglobin saturation brought about by a hyperoxic gas challenge^[2, 3]^. This measurement can be used to estimate the Cerebral Metabolic Rate of Oxygen Consumption (CMRO_2_)^[4]^, and can even quantify regional venous oxygenation in the brain^[1]^. To date however, research has been carried out exclusively in the cerebral vasculature. In this study, we aimed to explore whether this technique can be extended to the liver to noninvasively quantify hepatic venous oxygen saturation (ShvO_2_).

ShvO_2_ is an indicator of the oxygen supply to demand ratio in the liver^[7]^, and can currently only be measured invasively, via catheterisation. Several studies have shown the benefit of ShvO_2_ measurements^[8-10]^, particularly in patients that have undergone surgical procedures^[11]^. For example, it was found that after Fontan operations (a palliative procedure performed on children), monitoring ShvO_2_ in the immediate post-operative period could predict the occurrence and severity of subsequent acute liver dysfunction^[12]^. Likewise, it was shown that ShvO_2_ could be used to gauge the regeneration status of the remnant portion of the liver in rats that had undergone partial hepatectomy ^[11, 13]^.

Given the demonstrated benefits, we sought to investigate the potential of QSM to measure ShvO_2_ non-invasively. A hyperoxic gas challenge was administered to a cohort of healthy mice, which resulted in a controlled change in blood deoxyhaemoglobin content. Susceptibility was modelled in a large branch of the hepatic and portal veins and used to calculate SvO_2_ under normal and hyperoxic conditions. These oxygenation estimates were compared to gold-standard measurements with a blood gas analyser. In addition to this, SvO_2_ in mice that had been inoculated with colorectal liver metastases (CRLM) was also calculated in the portal and hepatic veins under normoxic conditions, and compared to that of the healthy mice. Given the increased metabolic burden that cancer places on host tissue, it is hypothesised that CRLM would result in a lower ShvO_2_ than in healthy mice.

## Materials and Methods

### Animal preparation

All animal studies were performed in accordance with the UK Home Office Animals Science Procedures Act (1986) and UK National Cancer Research Institute (NCRI) guidelines [14]. CD1 mice (n = 10) (female 8 – 12 weeks) were anaesthetised using 4% isoflurane in 100% O_2_. Respiratory rate was constantly monitored using a pressure pad (SA instruments, Stony Brook, NY USA) and maintained at ∼40 - 80 breaths per minute by varying isoflurane concentration between 1.5 and 3%. Body temperature was maintained at 37.5 ± 0.5^o^C using a warm water circulation system.

Colorectal liver metastases were induced in severe combined immunodeficiency (SCID, CD1 background) mice (n = 10), which were inoculated with 1x10^6^ SW1222 CRLM cells via intrasplenic injection^[15]^, followed immediately by a splenectomy. Mice were scanned at 19 days post-surgery.

Gasses were administered through a nose cone at a rate of 0.5 ltr/min. Images were acquired in all cases under normoxic conditions as the subject was administered medical air (21% O2/balance Nitrogen). Hyperoxia was induced in the healthy cohort via the administration of 100% O_2_. 10 minutes were allowed between gasses to allow the animals to acclimatise.

### MRI data acquisition

All subjects were scanned on a 9.4T MRI scanner (Agilent Technologies, Santa Clara, CA, USA) with a 39-mm-diameter birdcage coil (RAPID Biomed, Rimpar, Germany). Susceptibility data were acquired using single echo, T_2_^*^-weighted GRE acquisitions. The acquisition sequence was modified such that first order flow was compensated for in the x, y, and z, directions. Scan parameters were as follows: repetition time (TR) = 1000 ms; echo time (TE) = 4 ms; flip angle = 70^o^; voxel size = 200×200 µm^2^; readout acquisition bandwidth = 50 kHz; number of averages = 8, slice thickness = 200 µm^2^.

The field of view in each case was adjusted to ensure full coverage of the liver and water reference, and matrix size was adjusted such that voxel size was maintained and the data were spatially isotropic. The number of slices varied from 60 - 80 to accommodate full coverage of the liver. Mice bearing tumours were at an advanced stage of disease and did not tolerate anaesthetic well. As such, the number of signal averages was reduced to 4 to decrease scan time and ensure all mice survived the imaging protocol.

All MRI acquisitions were respiratory-gated. This was done by monitoring the animals breathing rate with a pressure sensitive respiratory monitor (SA Instruments, Stony Brook, NY, USA) while in the scanner. Data would only be acquired during a flat region of the respiratory cycle (i.e. between breaths) in order to avoid respiratory related motion artefacts in the image. Total scan time was 20 – 40 mins, depending on respiration rate.

### Water reference

For absolute quantification of susceptibility from QSM data, a reference must material must be used for calibration ^[16]^. In the brain, this is usually cerebrospinal fluid (CSF) within ventricles ^[17]^, but in the liver no comparable material is available. We therefore proposed the inclusion of a sample of distilled water in the scanner with each subject (figure 1). Briefly, a thin, cylindrical, nitrile membrane (∼6 cm length, ∼2 cm dia.) was filled with distilled water and sealed, with care taken to ensure no air became trapped in the process. This was placed beneath each mouse in the animal holder before scanning commenced. All susceptibility values are quoted with respect to the water reference.

**Fig 1:**
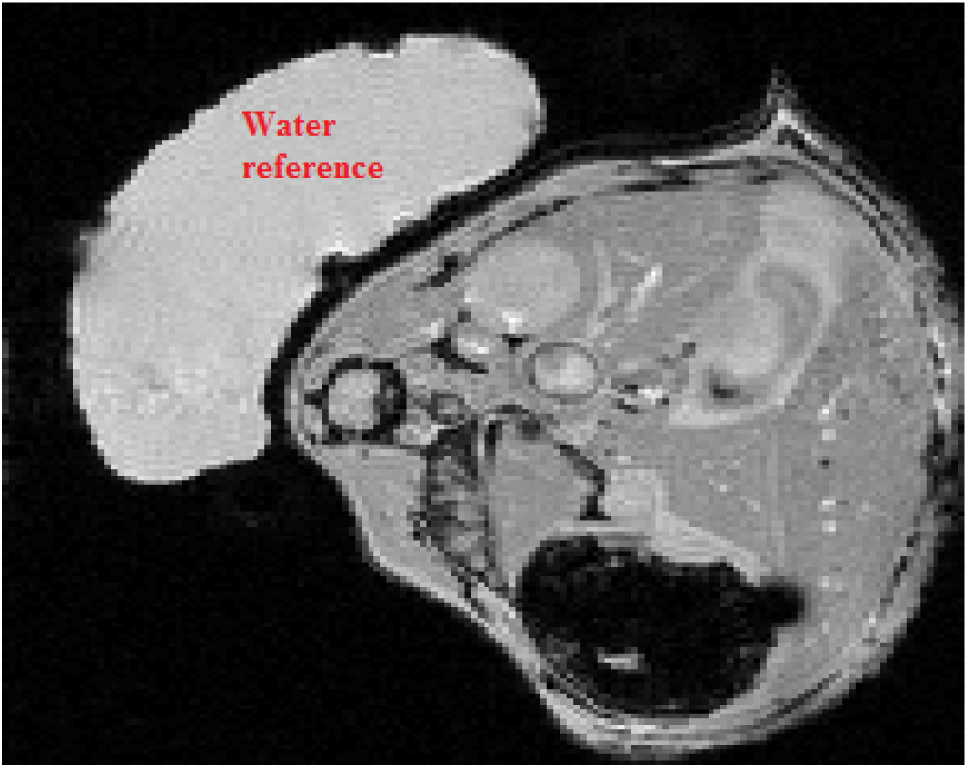
An example magnitude image of a mouse liver (axial orientation), showing the location of the water reference used to calibrate susceptibility measurements

### Image processing and analysis

Quantitative susceptibility maps were calculated from raw phase data. A binary mask was manually segmented around the entire liver in each magnitude image using ITK-SNAP ^[18]^. Phase unwrapping and background field suppression were performed using a Laplacian based SHARP algorithm (TSVD threshold = 0.04, mask erode = 2 – 3 voxels) ^[19]^.

Susceptibility inversion was carried out using the Thresholded K-Space Division (TKD) algorithm ^[20]^. The threshold of the TKD kernel was set to ± 0.2, such that absolute values outside of this range were set to the threshold value with the appropriate sign depending on the position of the voxel within the kernel. A correction factor of 1.26 (i.e. 1 / 0.786) was included in the deconvolution operation in the TKD algorithm, as recommended ^[19]^. All post processing was performed in Matlab (version 2015b, The MathWorks, Natick, MA).

Regions of interest (ROIs) were manually segmented on each magnitude image using ITK-SNAP ^[18]^, and corresponded to large branches of the hepatic vein (HV) and the portal vein (PV). In order to avoid partial volume effects, only voxels in the highest 20^th^ percentile of susceptibility values per ROI were accepted ^[1]^.

### Calculating ShvO_2_

The susceptibility difference between blood and water can be related to SvO_2_ by the following:

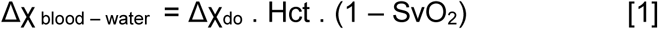

where ΔX_do_ = 2.26 ppm (SI) is the difference in susceptibility between fully oxygenated and deoxygenated blood ^[21]^, and Hct is the fraction of blood composed of haematocrit, assumed here to be 0.4 ^[3]^. The study cited relates SvO_2_ to the susceptibility shift between the vein and surrounding tissue. All calculations performed in this study used the water sample as the reference point.

### Blood gas measurement

As a gold standard measurement for comparison with QSM measurements, blood gases were measured invasively in three mice. Mice were anaesthetised as described above and a syringe was used to extract blood from a portion of the inferior vena cava (IVC) within the liver under ultrasound guidance. The procedure was carried out under normoxic and hyperoxic conditions for each mouse, by administering gases as per MRI experiments. 10 minutes were allowed following a change in administered gas to allow the animal to acclimatise before sampling. Samples were transferred from the syringe to a 2 µl heparinised glass tube, and then to the blood gas analyser (RAPIDLab 348EX blood gas system (Siemens)).

### Statistical analysis

Parameter estimates were compared using a Wilcoxon matched-pairs signed rank test, in which a difference was considered statistically significant for P < 0.05.

## Results

The susceptibility of the blood in the large branches of the liver vasculature became more diamagnetic in response to the administration of O_2_. This is illustrated in figure 2, which features maximum intensity projections of a segment of a susceptibility map of a healthy mouse liver (11 slices, 2.2 mm segment), calculated from data acquired under norm- and hyperoxic conditions. The blood vessels in the normoxic image are more prominent with respect to the liver tissue due to the increased presence of deoxyhaemoglobin.

**Fig 2.**
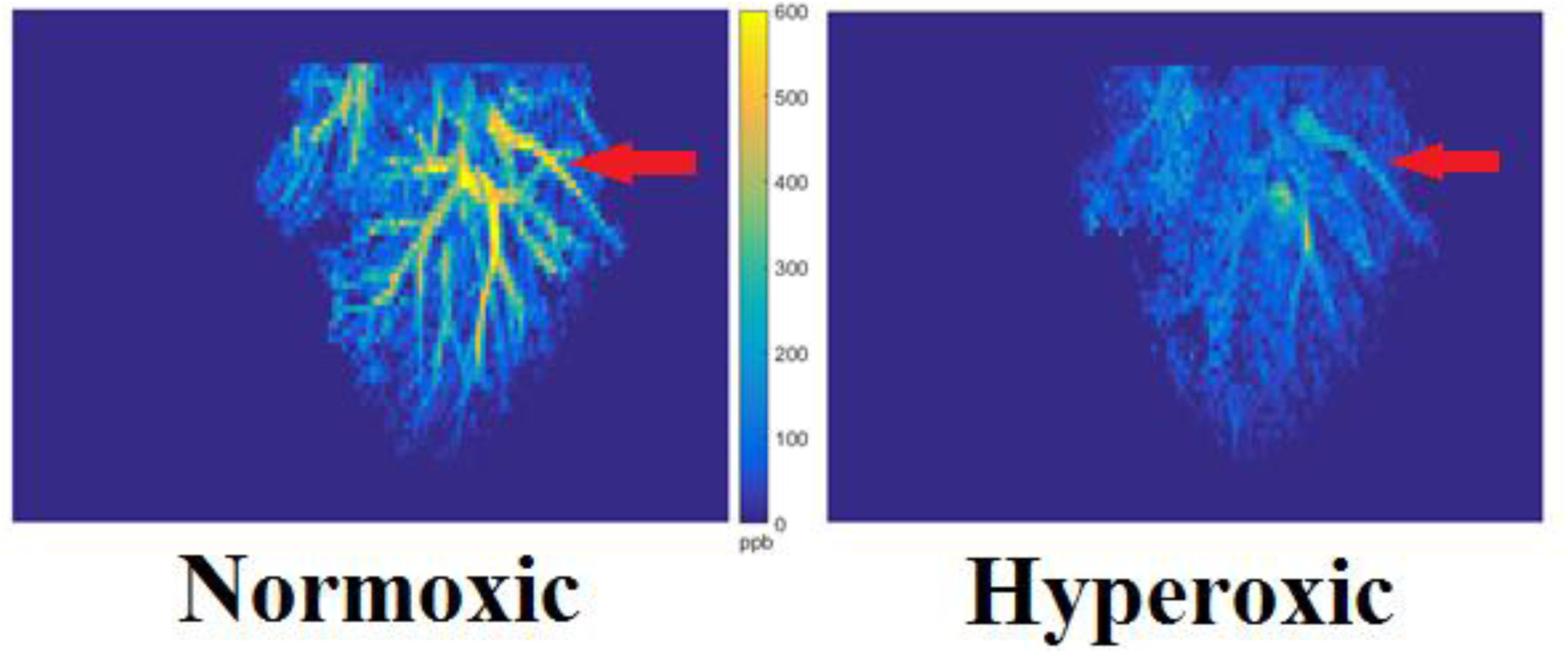
Maximum intensity projections of processed QSM data, from a 2.2 mm segment of a representative mouse liver, under normoxic and hyperoxic conditions. Large branches of the hepatic vein are clearly visible in each image (red arrows). Vessels are brighter with respect to the liver tissue (by approximately 500 ppb) in the normoxic image compared with the hyperoxic image, indicating a more paramagnetic susceptibility. This is due to the increased concentration of deoxyhaemoglobin in the blood under normoxia.

Magnetic susceptibility decreased significantly in the portal and hepatic veins of the healthy animals in response to hyperoxia (results summarised in table 1). Mean susceptibility of the blood in the portal vein decreased from 383 ± 134 ppb under normoxia to 248 ± 161 ppb under hyperoxia (p < 0.05 n = 10). Similarly, susceptibility decreased from 427 ± 161 ppb to 234 ± 80 ppb in the hepatic vein under normoxic and hyperoxic conditions respectively (p < 0.01, n = 10).

**Table 1:**
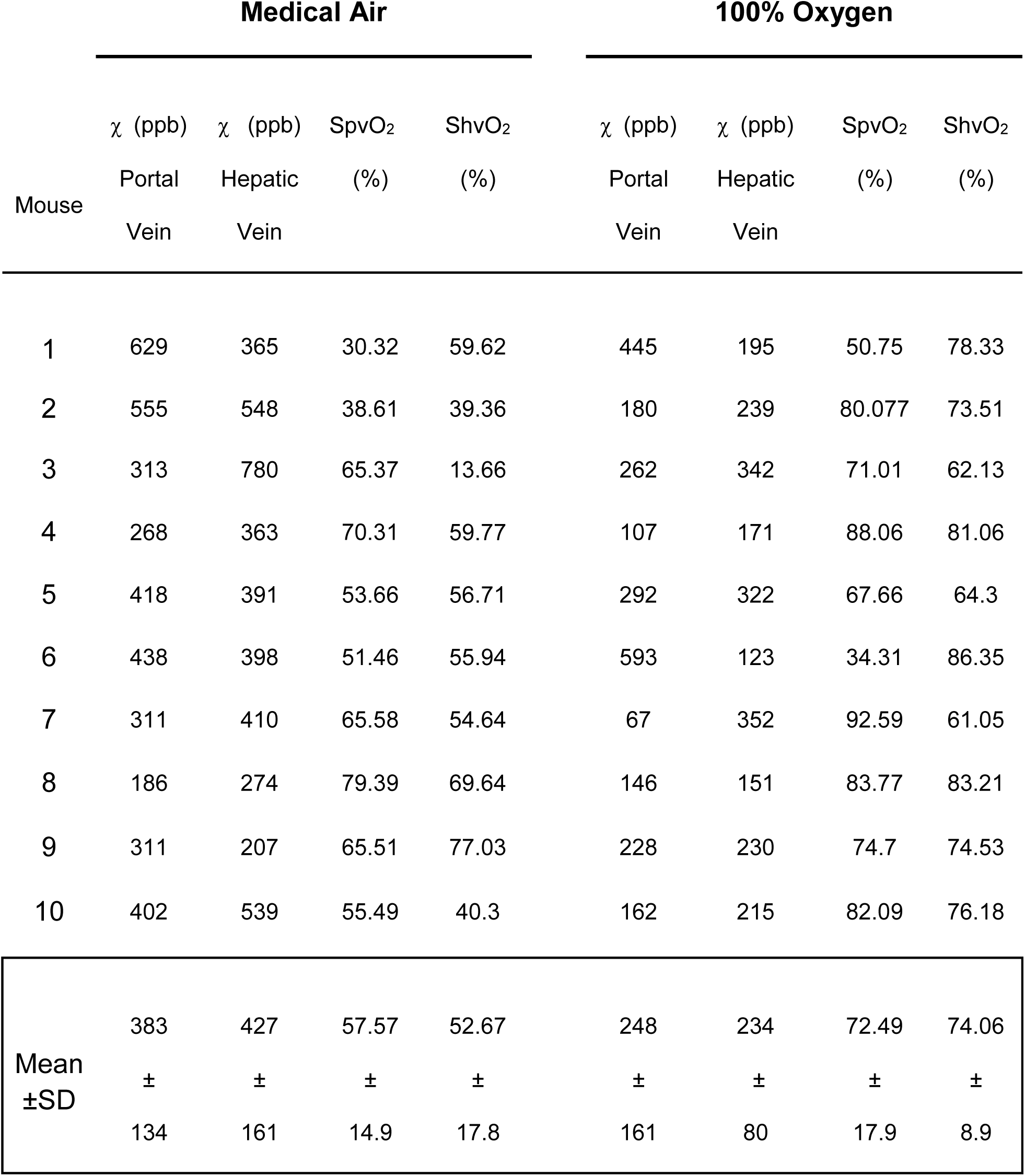
Magnetic susceptibility and venous oxygen saturation, measured non-invasively in portal and hepatic veins, during administration of medical air or 100% oxygen.

Venous oxygen saturation increased significantly in both the portal and hepatic veins of the healthy animals in response to the administration of pure O_2_ (fig. 3). In the portal vein (fig. 3A), SpvO_2_ increased by 14.93% from 57.57 ± 14.9% during air-breathing to 72.5 ± 17.9% during hyperoxia (p < 0.05, n = 10), while in the hepatic vein (fig. 3B), ShvO_2_ increased by 21.39% from 52.67 ± 17.8% to 74.06 ± 8.9% (p < 0.01). Results are summarised in table 1.

**Fig 3:**
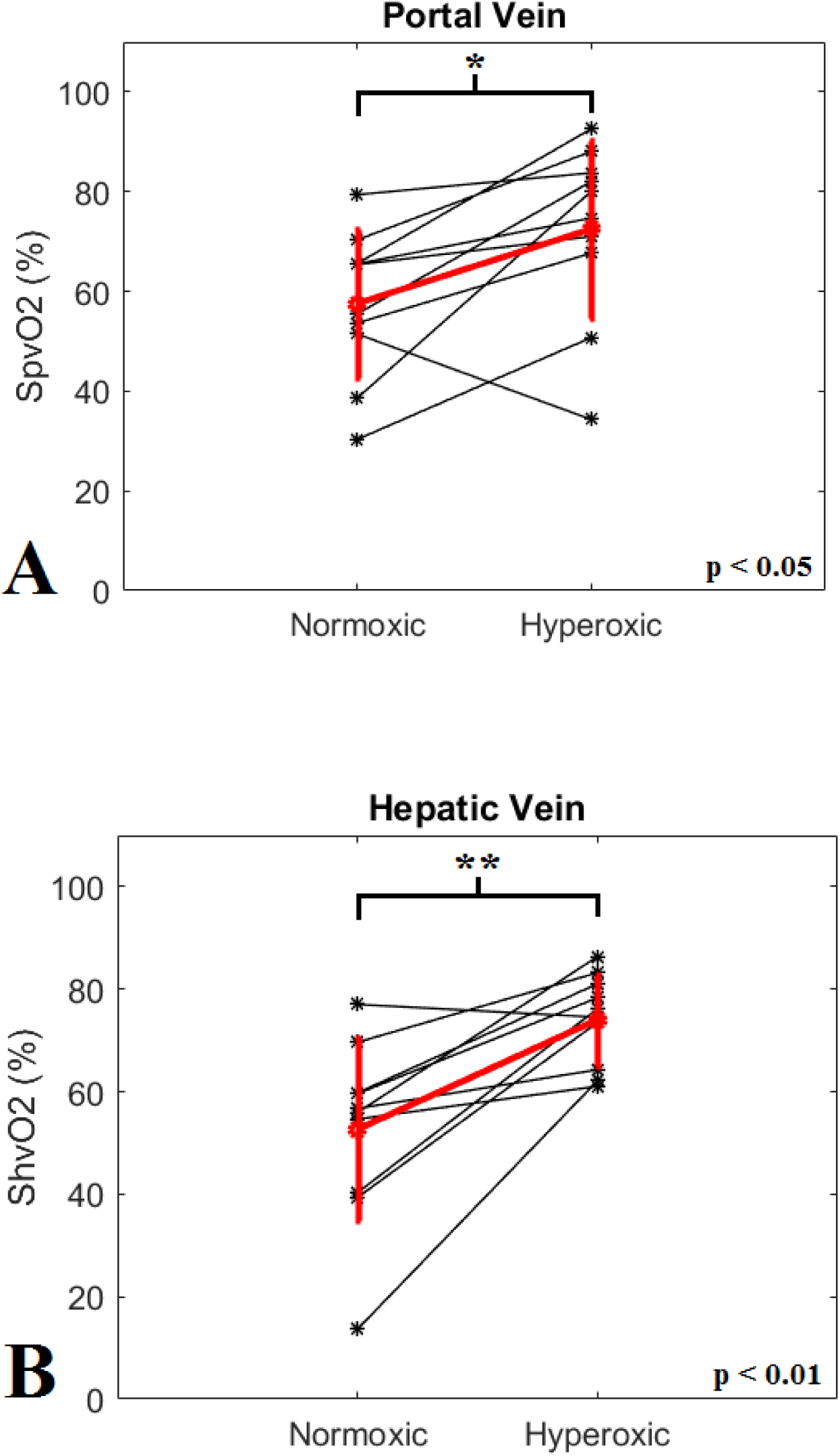
Graphs showing the change in hepatic venous oxygen saturation (ShvO_2_) in (a) the hepatic and (b) portal veins. A statistically significant increase in oxygen saturation was measured in response to hyperoxia in both vessels (*p<0.05,**p<0.01)

There was excellent agreement between the measurements of ShvO_2_ acquired on the blood gas analyser and that calculated from QSM measurements (fig. 4). The invasive measurement showed a mean increase of 26.14% in blood oxygenation, from 52.83 ± 9.78% to 78.97 ± 11.45%. The difference was not statistically significant.

**Fig 4:**
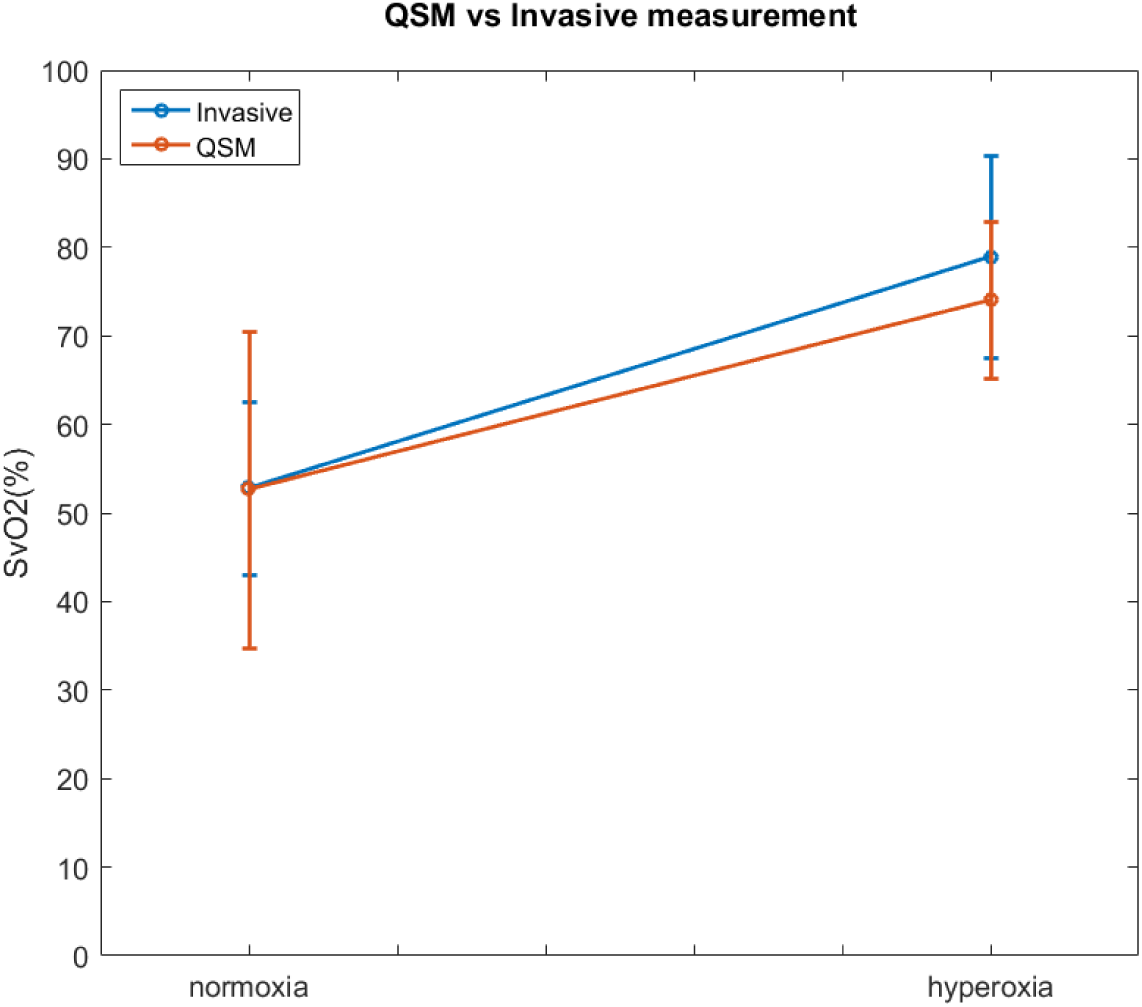
Graph showing the change in venous oxygen saturation in the hepatic vein from: non-invasive measurements with QSM; and invasive measurements from the IVC with a blood gas analyser. There is an excellent agreement between the two datasets.

The graphs in figure 5 show the venous blood oxygen saturation measured under normoxic conditions in the portal (A) and hepatic (B) veins of the mice with tumours and the healthy cohort. Mean oxygen saturation in the portal vein was 43.84 ± 23.1% and 57.57 ± 14.9% for the disease and healthy animals respectively, and in the hepatic vein was 30.18 ± 11.6% and 52.67 ± 17.8% for the disease and healthy animals respectively. There was no significant difference between the cohorts measured in the portal vein, however the oxygen saturation of the blood in the hepatic vein was significantly lower in the mice with tumours, when compared to the wild types (p < 0.05). As the effect was not observed in the portal vein, this would indicate that the effect is not systemic, but is instead caused as the blood passes through the liver. It is expected that this can be attributed to the increased metabolic requirements of the liver tumours.

**Fig 5:**
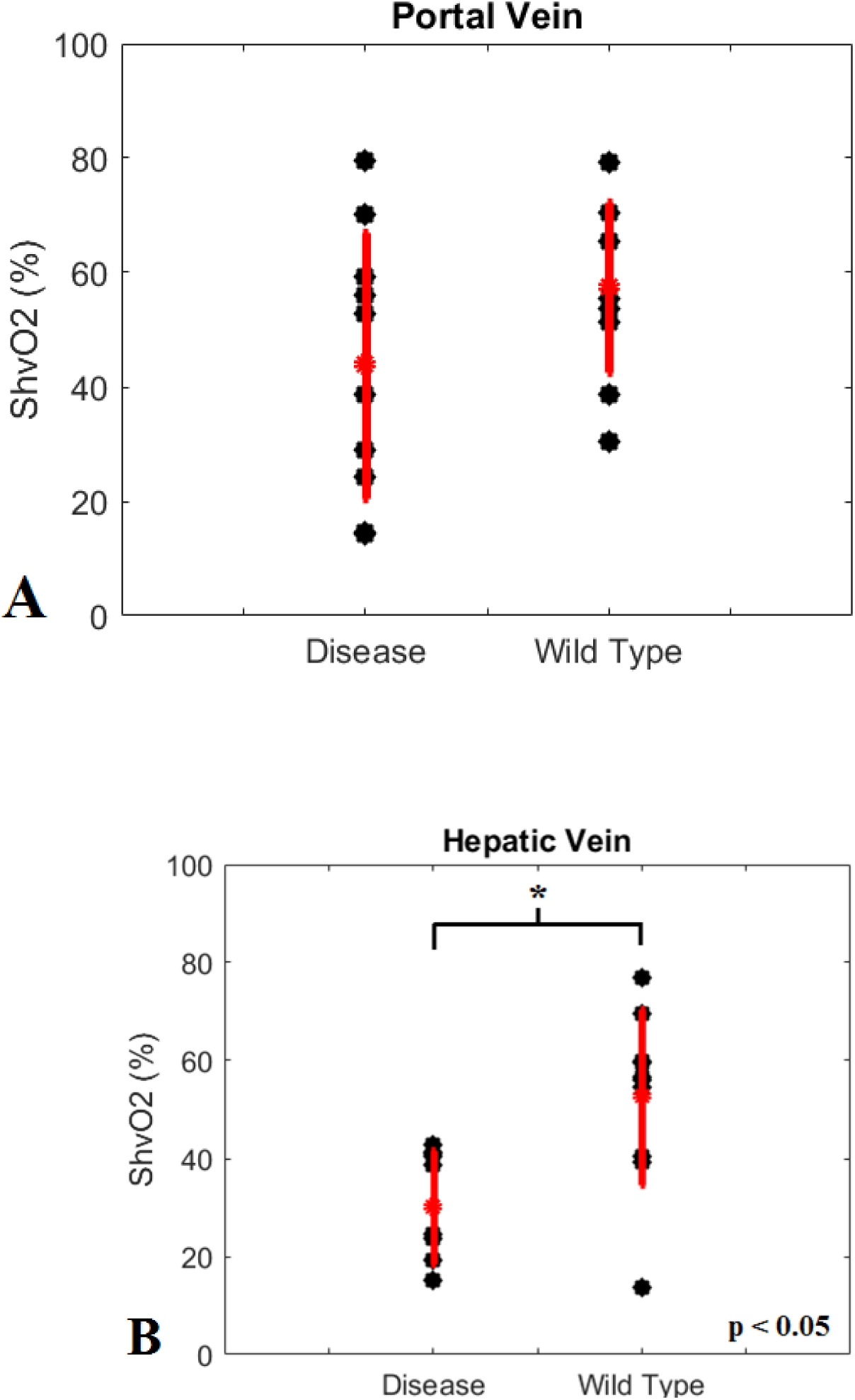
Graphs showing measurements of venous blood oxygen saturation in mice with tumours and healthy wild type, calculated from susceptibility measurements in the portal vein (A) and the hepatic vein (B). Measurements in the hepatic vein of the mice with tumours contained significantly less oxygen than the healthy cohort., presumably due to the increased metabolic demands of the tumour tissue.

## Discussion

In the current study, we employed QSM to assess changes in ShvO_2_, which we modulated with a hyperoxic gas challenge. This is the first report that examines the ability of QSM to assess oxygen changes in the major hepatic vessels, and we have shown that it is possible to detect statistically significant differences in blood oxygenation in response to hyperoxia. Moreover, our measurements showed a good accordance with invasive, gold-standard measurements with a blood gas analyser, and that it is possible to detect significant differences between the hepatic venous oxygen saturation of a group of healthy animals and a group with liver cancer.

A previous study that calculated SvO_2_ from QSM measurements in the cerebral vasculature of mice have reported changes from ∼88% to ∼99% in response to hyperoxia ^[3]^, which is an increase of roughly 10%. Absolute hepatic ShvO_2_ values calculated in the current study were much lower (approximately 55% during normoxia to 74% during hyperoxia), but can be attributed to blood having passed through the gut and mesentery before reaching the liver, and so it is expected that values would be lower than in the brain.

Previous reports in the literature of the response of portal venous blood in rats to hyperoxia describe an increase from ∼53% under normoxia to ∼93% under hyperoxia in healthy control animals ^[22]^. It is highly encouraging that this nomoxic SpvO_2_ value is comparable with the measurement made in our experiment. Indeed, whilst the increase under induced hyperoxia was greater than we have observed, this could be due to differences between experimental protocols, the efficiency of oxygen delivery and differences between species.

Moreover, a good agreement was found between QSM measurements and invasive blood gas measurements, during both normoxia and hyperoxia. The collection of murine hepatic blood is technically challenging due to the size of the vessels under examination and the small volume of blood that can be sampled in a mouse. On this basis, extraction of blood from the portal vein was not possible, and comparisons were instead drawn from measurements in the IVC, which was assumed to be representative of measurements in the smaller vessels.

The difference in hepatic venous blood saturation between the mice with tumours and healthy wild type mice could have significant clinical potential. The data in both cases were acquired under normoxic conditions so their acquisition necessitated little more than a single standard T_2_^*^-weighted scan. Future experimental work could be to perform a longitudinal study in order to characterise the correlation between tumour burden and ShvO_2_. Once this is established, it opens the possibility of using QSM to non-invasively diagnose or monitor liver cancer, differentiate between benign and malignant lesions, or even to gauge the efficacy of treatment regimes.

One of the limitations of this study was the inability to measure the susceptibility of the hepatic artery. At the resolution of the imaging protocol used here the diameter of the hepatic artery is of the order of a single voxel, so measurements were undermined by partial volume effects which are known to result in inaccurate estimations of susceptibility (ref). The ability to measure the susceptibility, and subsequently calculate the oxygen saturation of all three major hepatic vessels would allow a more complete characterisation of hepatic haemodynamics, as well as giving greater insight into hepatic oxygen metabolism, increasing the clinical usefulness of the technique. Hepatic arterial susceptibility measurements may be made possible by acquiring higher resolution images, or by using larger animals (i.e. rats), however difficulty may be encountered due to pulsatile flow in the arterial vessel.

From a clinical translational point of view, this experiment should be relatively easy to implement in humans. It has been shown in a recent study that QSM data from the entire liver can be acquired in ∼19 seconds, i.e. within one breath hold ^[23]^. Data could simply be acquired under normoxic conditions, but equally, both patients and healthy volunteers can tolerate hyperoxia well. Previous experiments examining the use of QSM in the liver ^[24, 25]^ have focussed on quantifying liver iron, requiring complicated modifications of the acquisition and processing protocols in order to account for fat. One major advantage of measuring the susceptibility of blood in the large vessels is that no fat is present, and data can be acquired and processed via standard means.

Our use of an external susceptibility reference is a straightforward solution for absolute susceptibility calibration outside the brain. Susceptibility references are required to be independent of experimental variables, easy to depict and delineate, and easily identifiable across a wide range of subjects ^[17]^. The external reference used here meets each of these criteria. The inclusion of a reference phantom in the experimental setup is straightforward, although one limitation is that the image field of view must be increased to accommodate it, potentially resulting in longer acquisition times if resolution is to be maintained.

The ability to non-invasively perform venous oximetry in the liver could have important clinical implications. Hepatic venous oxygen saturation is an eminently useful metric, and has been used to assess hepatic oxygen kinetics in studies with respective focusses as diverse as haemodialysis ^[26]^, acute and chronic heart failure ^[27]^, and hepatic ischemic/reperfusion injuries ^[28, 29]^. Furthermore, thanks to improvements in diagnostic radiology, patient selection and operative technique, partial hepatectomy has increasingly become a more viable treatment option in cases of hepatic lesions, both malignant and benign. It is known that the regenerating liver places an increased metabolic burden on patients that have undergone the procedure, and it has been shown previously that ShvO_2_ reflects the metabolic status of the remnant liver ^[11, 13]^. The advent of QSM means that the ability to relate magnetic susceptibility to ShvO_2_ offers a way to asses this in a non-invasive and repeatable fashion.

